# Krüppel like factor 2 – deficient myeloid cells promote skeletal muscle regeneration after injury

**DOI:** 10.1101/312868

**Authors:** Palanikumar Manoharan, Taejeong Song, Tatiana L Radzyukevich, Sakthivel Sadayappan, Jerry B Lingrel, Judith A Heiny

## Abstract

Regeneration of adult skeletal muscle after injury is coordinated by complex interactions between the injured muscle and the innate immune system. Myeloid lineage cells predominate in this process. This study examined the role of Krüppel – like factor 2 (KLF2), a zinc-finger transcription factor that regulates myeloid cell activation state, in muscle regeneration. Gastrocnemius muscles of wild-type and *myeKlf2^-/-^* mice, which lack KLF2 in all myeloid cells, were subjected to cardiotoxin injury and followed for 21 days. Injured muscles of *myeKlf2^-/-^* contained more infiltrating, inflammatory Ly6C^+^ monocytes, with elevated expression of inflammatory mediators. Infiltrating monocytes matured earlier into pro-inflammatory macrophages with phenotype Ly6C^+^, CD11b^+^, F4/80^+^. Inflammation resolved earlier and progressed to myogenesis, marked by an earlier decline of Ly6C^+^ macrophages and their replacement with anti-inflammatory Ly6C^-^ populations, in association with elevated expression of factors that resolve inflammation and promote myogenesis. Overall, regeneration was completed earlier. These findings identify myeloid KLF2 as a central regulator of the innate immune response to acute skeletal muscle injury. Manipulating myeloid KLF2 levels may be a useful strategy for accelerating regeneration.

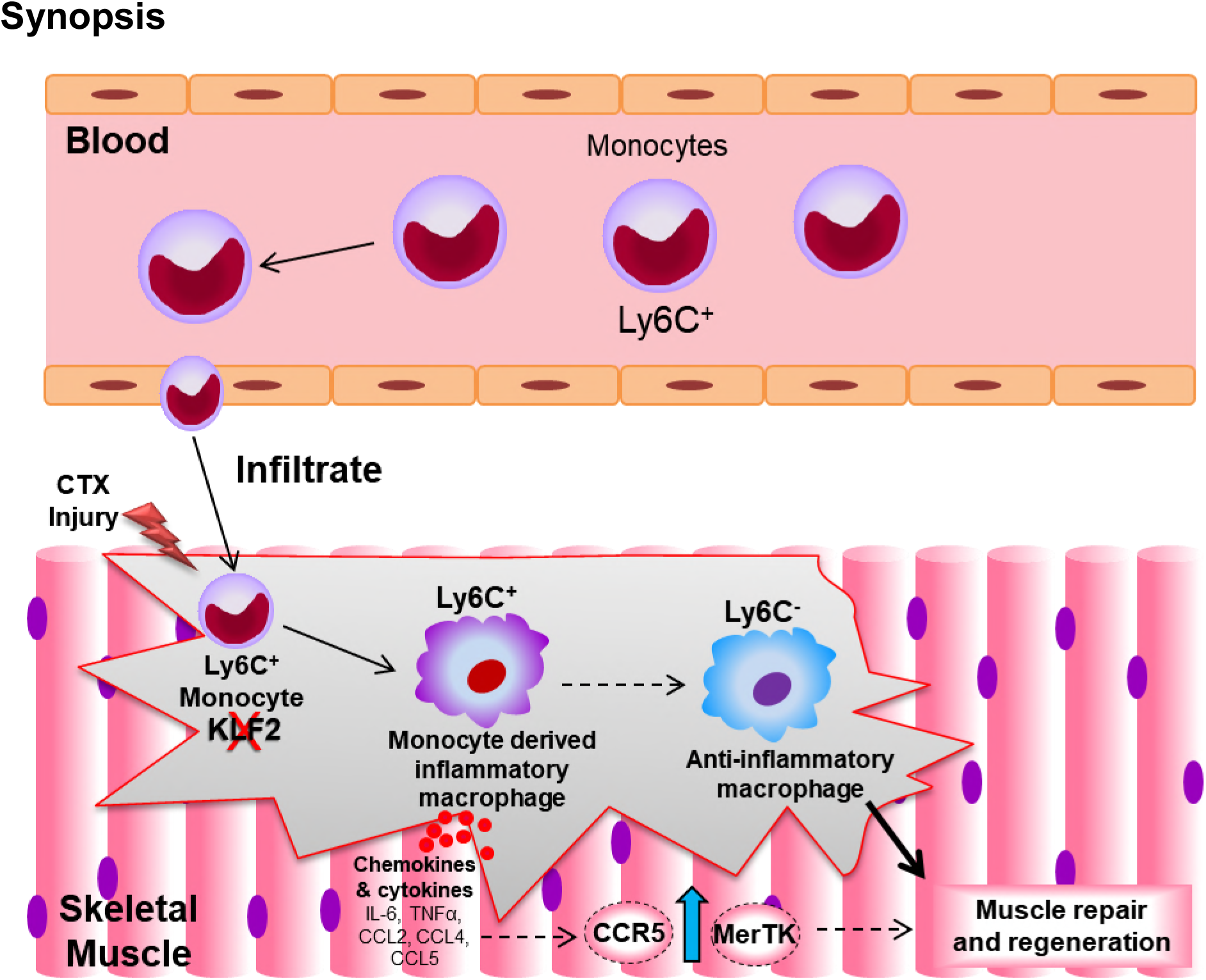
Synopsis. The zinc-finger transcription factor, KLF2, is a central regulator of the innate immune response to skeletal muscle injury. Targeted deletion of KLF2 in myeloid lineage cells of mice (*myeKlf2^-/-^*) enhances the immune response and, notably, accelerates muscle repair.

- Injured muscles of *myeKlf2^-/-^* mice recruit greater numbers of Ly6C^+^ (inflammatory) monocytes from the circulation.
- These mature in situ into pro-inflammatory macrophages which phagocytose necrotic tissue and prepare the environment for muscle regeneration.
- Subsequently, Ly6C^+^ macrophages decline and are replaced by anti-inflammatory (Ly6C^-^) macrophages that promote myogenesis of new fibers.
- Injured muscles of *myeKlf2^-/-^* mice complete regeneration earlier, with phenotypically adult fibers.

## Introduction

Adult skeletal muscles retain a remarkable capacity to regenerate lost or damaged tissue. Skeletal muscle regeneration after acute injury progress through two main, overlapping phases, broadly defined as an initial proinflammatory phase followed by an anti-inflammatory, regenerative phase [1-5]. The regenerative phase recapitulates in many ways the process of embryonic myogenesis, while the immune response is unique to adult skeletal muscles. The immune response is essential for proper completion of repair [6].

Muscle regeneration after injury is orchestrated by coordinated interactions between skeletal muscle and the innate immune system. Myeloid lineage immune cells play a central role in this process [4, 5, 7-9]. Indeed, depletion of circulating monocytes slows regeneration and reconstitution of myeloid lineage cells restores regeneration [10]. Myeloid lineage monocytes dominate the initial inflammatory infiltrate, and are present at up to 100,000 cells/mm^3^ of muscle [11]. As regeneration progresses, myeloid cells assume a continuum of activated phenotypes that support each phase of repair [7, 12]. Macrophages remove necrotic tissue, secrete cytokines, chemokines, and growth factors, induce production of the fibrotic scaffold and blood supply needed for the growth and attachment of new fibers, help resolve the inflammation, and support the growth and differentiation of new myofibers [5, 13]. Macrophages eventually disappear as regeneration is completed and newly formed fibers with adult phenotypes are integrated into the muscle syncytium. The molecular mechanisms and signaling molecules that determine myeloid lineage phenotypes and functions during skeletal muscle repair remain poorly understood.

The zinc-finger transcription factor, KLF2, plays a central role in the activation and phenotypic determination of a variety of immune cell types, both lymphoid and myeloid [14-16]. KLF2 is a central, negative regulator of gene clusters which mediate the proinflammatory activation of monocytes and macrophages *in vitro* [17], and its loss promotes an inflammatory monocyte/macrophage profile *in vitro* and *in vivo* [15, 18, 19]. Macrophages of mice which lack KLF2 in all myeloid cells assume an enhanced inflammatory profile, with elevated expression of inflammatory mediators and greater infiltration into atherosclerotic plaques [18, 19].

Considering the central role of myeloid lineage cells in skeletal muscle repair and the role of KLF2 in determining their activation state, we hypothesized that KLF2 may play an important role in the immune response to acute muscle injury. To test this hypothesis, we compared regeneration in injured skeletal muscles of WT and *myeKlf2*^-/-^ mice. We subjected the gastrocnemius muscle to cardiotoxin (CTX) or sham injury and followed regeneration for 21 days post injury. The progress of regeneration was assessed from the expression levels of inflammatory mediators and myogenic markers, macrophage phenotypes, and histological changes.

Our results show that loss of myeloid KLF2 significantly enhances the inflammatory immune response to muscle injury and, notably, that an enhanced inflammatory response accelerates regeneration. These findings demonstrate that myeloid KLF2 plays a central role in orchestrating the immune response to acute muscle injury. In addition, because myeloid cells are accessible from the circulation, manipulating KLF2 in myeloid lineage cells may be a useful therapeutic strategy for immune-based therapies to improve outcomes after muscle injury.

## Results

### Loss of KLF2 in myeloid-derived cells enhances the inflammatory immune response to muscle injury

The proinflammatory phase of muscle regeneration after acute injury is marked by significantly increased levels of inflammatory mediators, many of which are produced within the muscle by myeloid-derived monocytes and macrophages. To investigate the role of myeloid KLF2 in their production, we compared expression of the proinflammatory mediators — *Cox2*, *IL-6*, *Ccl2*, *Ccl4*, *Ccl5*, and *Tnfα* — in regenerating gastrocnemius muscles of WT and *myeKlf2*^-/-^ mice at days 3 and 5 post-CTX or -vehicle treatment (Fig. 1). This was found to be the period when the greatest differences between WT and *myeKlf2^-/-^* were seen.

**Figure 1.**
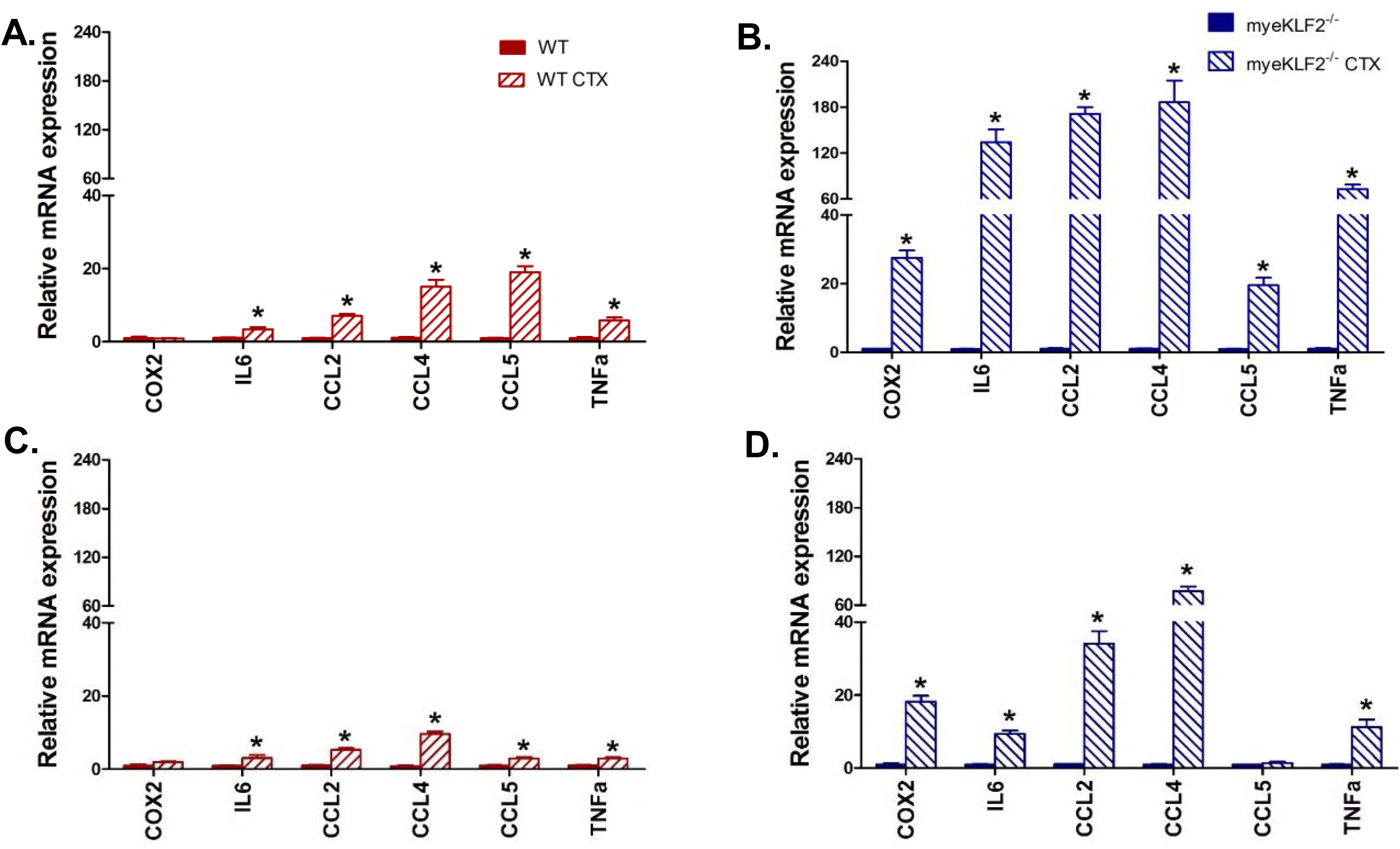
Loss of KLF2 in myeloid-derived cells enhances the inflammatory immune response to muscle injury. **A & B.** RT-qPCR analysis for inflammatory markers in gastrocnemius muscles of WT (red) and *myeKlf2^-/-^* (blue) mice were taken at day 3 following placebo (solid fill) or CTX (striped) injury. **C & D.** RT-qPCR analysis for inflammatory markers in gastrocnemius muscles of WT (red) and *myeKlf2^-/-^* (blue) mice were taken at day 5 following placebo or CTX injury. ^*^ indicates significant difference at p < 0.05; n = 4.

Each of these cytokines/chemokines plays a known role in inflammation and regeneration. IL-6 is one of the earliest cytokines detected after acute muscle injury and the majority of IL-6 is produced by monocytes recruited from the circulation [20]. The signaling cytokines, *Ccl2*, *Ccl4*, and *Ccl5*, are released from the injured tissue and are also secreted by infiltrating monocytes. *Ccl2 is* regulated by KLF2 via miR124a and miR150 [18]. The cytokines TNFα and COX2, which mediate inflammation through prostaglandins, originate from activated, macrophages that matured *in situ* from infiltrating monocytes. *Cox2* signaling is essential during the early stages of skeletal muscle regeneration [21].

In injured muscles of WT mice, *IL-6*, *Ccl2*, *Ccl4*, *Ccl5*, and *Tnfα* are present at day 3 post injury (Fig. 1A) and decline by day 5 (Fig.1C).Their appearance indicates that recruitment and infiltration of circulating monocytes is underway. Their decline by day 5 indicates that inflammation is resolving. Expression of *Tnfα* indicates that some of the infiltrating monocytes have matured into activated, proinflammatory macrophages. Injured muscles of *myeKlf2*^-/-^ mice express these markers at 15 to 30-fold greater levels than WT at day 3 post injury (Fig. 1B); all but *Ccl5* remain significantly elevated at day 5 (Fig 1D), a time when their expression has returned to near baseline in WT. None of these markers are significantly expressed in muscles of either genotype after placebo injury. The dramatically increased expression of inflammatory mediators and their earlier decline in muscles of *myeKlf2*^-/-^ mice suggest that loss of KLF2 in myeloid cells enhances and accelerates the proinflammatory phase of skeletal muscle regeneration after acute injury.

### Injured muscles of myeKlf2^-/-^ mice recruit greater numbers of circulating Ly6C^+^ monocytes than WT mice

The enhanced expression of inflammatory markers in injured muscles of *myeKlf2*^-/-^ mice suggests that these muscles recruit greater numbers of inflammatory Ly6C^+^ monocytes from the circulation. This population provides the pool that matures *in situ* into monocyte-derived macrophages. To examine this possibility, we compared the number of Ly6C^+^ cells present in regenerating skeletal muscles of *myeKlf2*^-/-^ and WT mice at day 3 (Fig. 2A, 2B) and day 5 (Fig. 2D, 2E) post injury. Ly6C^-^ cells are not present in uninjured skeletal muscles, and injured skeletal muscles do not recruit Ly6C^-^ monocytes [5].

**Figure 2.**
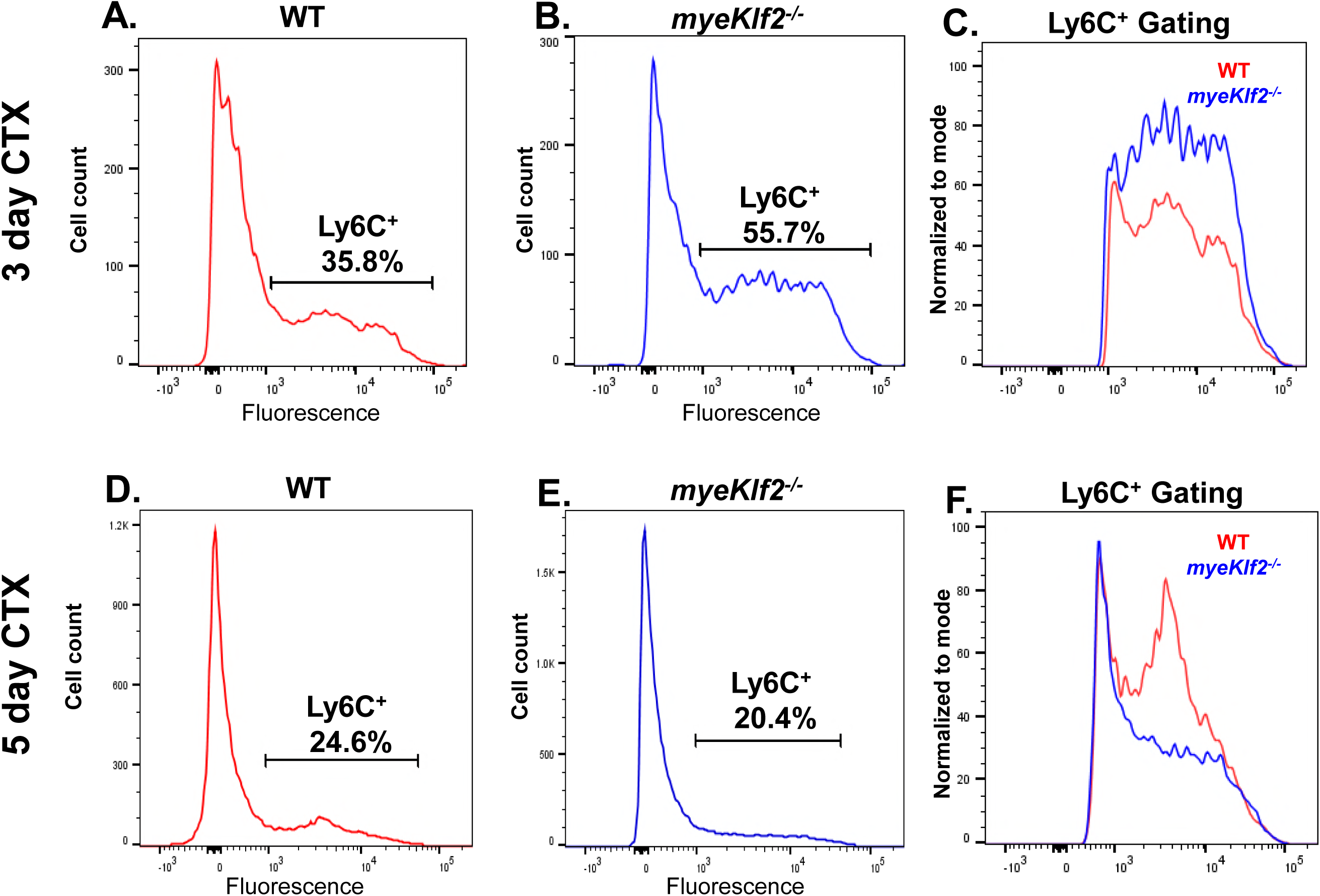
Injured muscles of *myeKlf2*^-/-^ mice recruit greater numbers of circulating Ly6C^+^ monocytes than WT mice. **A – C.** Representative flow cytometric analysis showing Ly6C^+^ cells present in injured muscles of WT (A) and *myeKlf2*^-/-^ (B) mice at day 3 following CTX injury. Vertical axis: Ly6C count. Horizontal axis: number of cells. Results are expressed as the percentage of mononuclear cells isolated from muscle. The panel 2C is the comparative graph of Ly6C^+^ expression between WT and *myeKlf2*^-/-^ mice at day 3 (n = 6; p<0.05). **D – F.** Representative flow cytometric analysis showing Ly6C^+^ cells present in injured muscles of WT (D) and *myeKlf2*^-/-^ (E) mice at day 5 following CTX injury. The panel 2F is the comparative graph of Ly6C^+^ expression between WT and *myeKlf2*^-/-^ mice at day 5 (n = 6; p<0.05).

Ly6C^+^ monocytes and macrophages are present in injured muscles of both genotypes. However, by day 3, injured muscles of *myeKlf2*^-/-^ mice have recruited a significantly greater number of Ly6C^+^ monocytes than WT (Fig. 2A, 2B, 55.7% vs. 35.7%, n=6; p <0.05). The number of Ly6C^+^ monocytes subsequently declines and this decrease occurs first in *myeKlf2*^-/-^ muscles. It is apparent by day 5, a time when their number is still increasing in WT (Fig. 2D, 2E, 20.4% vs. 24.6%, n=6; p <0.05). Overall, injured muscles of *myeKlf2*^-/-^ mice have recruited a significantly greater number of circulating Ly6C^+^ cells at day 3 (Fig. 2C, rightmost panels, blue vs. red counts) and this cell population declines earlier in injured muscles of *myeKlf2^-/-^* compared to WT mice (Fig. 2F).

### Infiltrating Ly6C^+^ monocytes mature earlier into inflammatory macrophages that are phenotypically Ly6C^+^, CD11b^+^, F4/80^+^

Proinflammatory macrophages within the injured tissue derive from circulating Ly6C^+^ monocytes [5]. Macrophages in regenerating skeletal muscles do not assume the canonical M1/M2 phenotypes defined for other inflammatory conditions [22, 23]. To identify the relevant populations, we further sorted immune cells by CD11b and F4/80 markers (Fig. 3). Myeloid lineage monocytes and macrophages express abundant surface marker CD11b, which is upregulated upon activation. F4/80^+^ is a macrophage-specific marker of monocyte differentiation that identifies intramuscular, mature macrophages. It is not found in circulating monocytes. Cells which are positive for all 3 markers – Ly6C^+^, CD11b^+^, and F4/80^+^ – represent the population of intramuscular, proinflammatory macrophages which derived from circulating Ly6C^+^ monocytes. Differences between WT and *myeKlf2*^-/-^ reveal populations that are influenced by KLF2.

**Figure 3.**
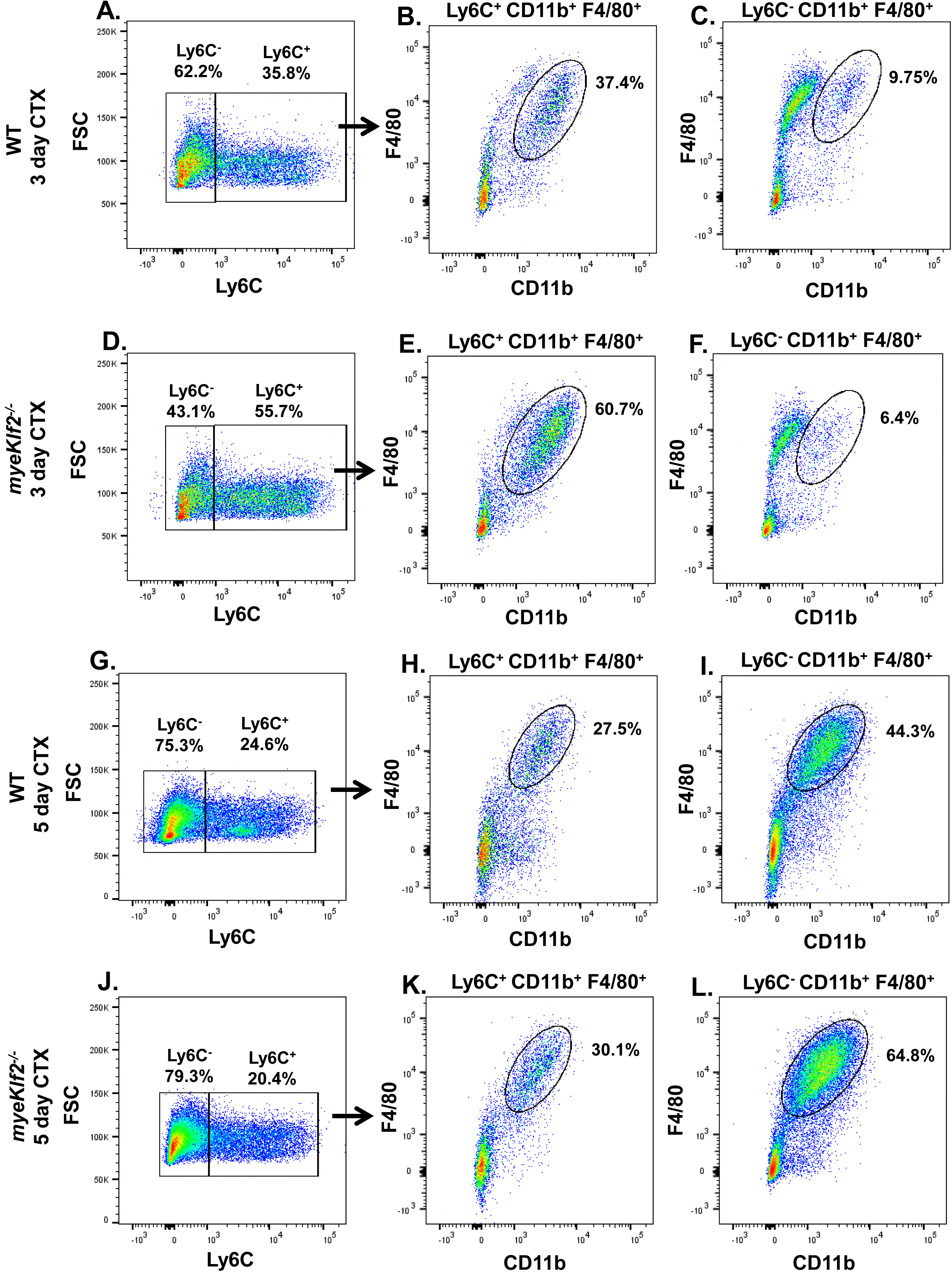
Infiltrating Ly6C^+^ monocytes mature earlier into inflammatory macrophages that are phenotypically Ly6C^+^, CD11b^+^, F4/80^+^. **A – F.** Day 3 post injury: Ly6C^+^ and Ly6C^-^ cells were gated from regenerating gastrocnemius muscles of WT (**3A to 3C**) and myeKlf2^-/-^ (**3D to 3F**) mice, and further sorted by flow cytometry using antibodies against CD11b (horizontal) and F4/80 (vertical). **3B** and **3E**, macrophages that are triple positive for Ly6C^+^, CD11b^+^, and F4/80^+^ (circled). **3C** and **3F**, macrophages that are triple positive for Ly6C^-^, CD11b^+^ and F4/80^+^ (n =6; p<0.05). **G – L.** Day 5 post injury: Same sorting strategy. **3G** to **3I**, WT; **3J** to **3L**, myeKlf2^-/-^. Circles indicate the triple positive cells used for counting (n =6; p<0.05).

By day 3, proinflammatory, monocyte-derived macrophages are evident in both genotypes. This population is significantly greater in *myeKlf2*^-/-^ compared to WT (Fig. 3E vs. 3B; 60.7% in *myeKlf2*^-/-^ vs. 37.4% in WT; n=6; p < 0.05). Only trace numbers of anti-inflammatory, Ly6C^-^ cells are detected in both genotypes at day 3 (Fig. 3C and 3F, 9.75% and 6.4%, n=6; p > 0.05). The near absence of Ly6C^-^ cells at day 3 is consistent with a previous report that injured skeletal muscles recruit only Ly6C^+^ monocytes from the circulation [5].

By day 5, the Ly6C^+^ population declines in both genotypes (Fig. 3H and 3K; *see also Fig. 2*) and the decline is associated with an increase in anti-inflammatory Ly6C^-^ populations (Fig. 3I and 3L). The majority of these cells are phenotypically Ly6C^-^, CD11b^+^, F4/80^+^. This population is significantly greater in *myeKlf2^-/-^* compared to WT (Fig. 3L vs. 3I: 64.8% in *myeKlf2*^-/-^ vs. 44.4% in WT, n=6; p <0.05). The decline in Ly6C^+^ macrophages and associated increase in Ly6C^-^ macrophages indicates that inflammation is resolving. The greater abundance of anti-inflammatory macrophages in *myeKlf2*^-/-^ at day 5 suggests that resolution of inflammation is advanced in *myeKlf2*^-/-^ compared to WT.

### Expression of CCR5 and MerTK is enhanced in injured muscles of myeKlf2^-/-^ mice

Collectively, the findings reported in Figs. 1 — 3 show that loss of KLF2 enhances the inflammatory phase of muscle regeneration after injury. To further define the role of KLF2 in this process, we compared the quantity of cells in each genotype expressing specific inflammatory and anti-inflammatory receptors (Fig. 4). KLF2 controls the expression of inflammatory chemokine receptors in other myeloid cells [16] and the set of expressed receptors determines, in part, the macrophage phenotype. For example, only activated, inflammatory T cells, which downregulate KLF2, express the chemokine receptor CCR5. Deletion of KLF2 in T cells induces expression of the CCR5 gene, which has consensus KLF2 binding sequences [16]. Interestingly, CCR5 is the cytokine receptor for CCL4 (MIP1β) and CCL5 (RANTES), both of which are dramatically elevated in injured muscles of *myeKlf2^-/-^* mice (Fig. 1).

**Figure 4.**
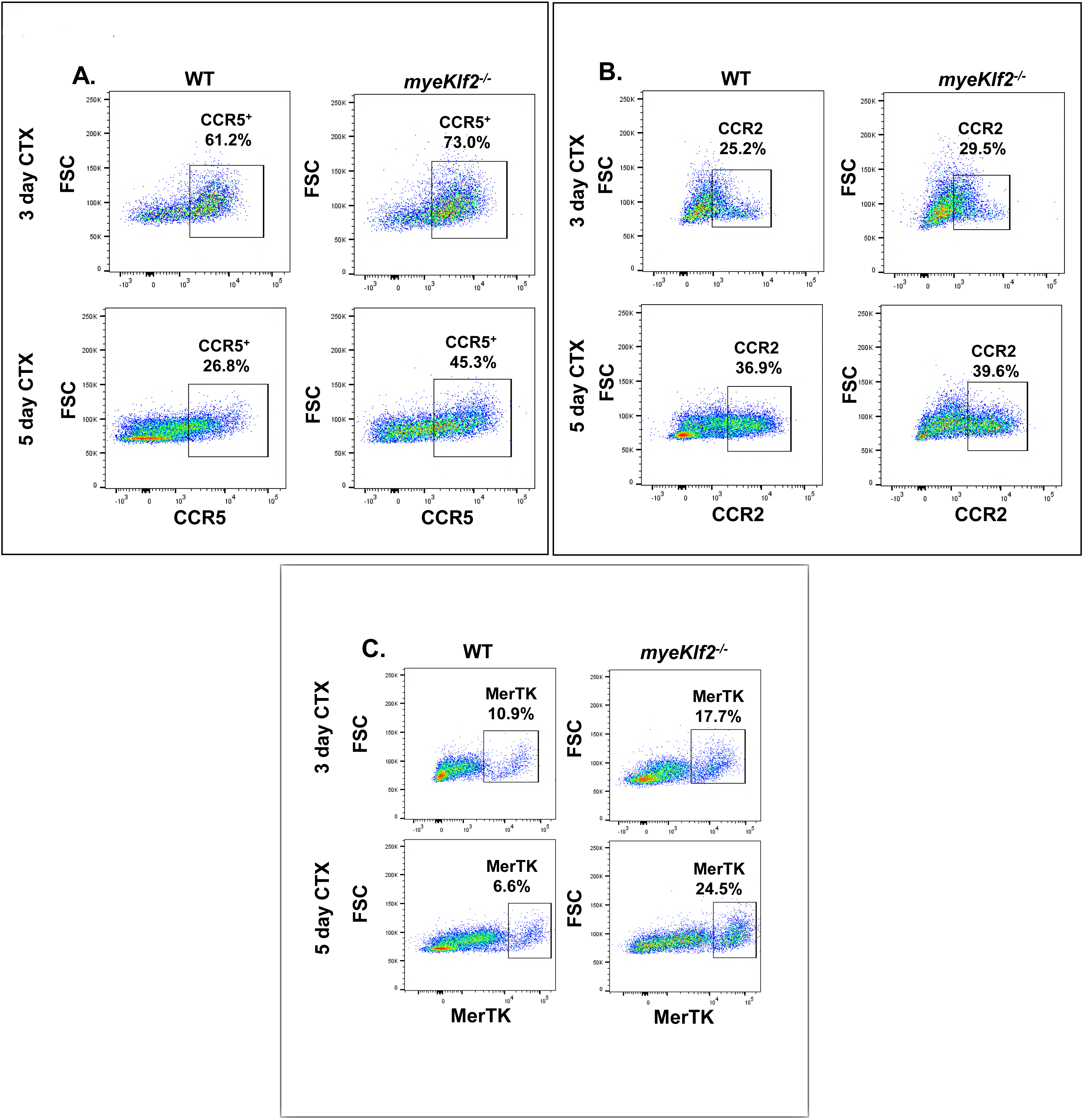
Expression of CCR5 and MerTK is enhanced in injured muscles of myeKlf2^-/-^ mice. **A.** Gastrocnemius muscles were obtained from WT and *myeKlf2^-/-^* mice at days 3 and 5 post CTX injury and stained and sorted by flow cytometry using antibody against CCR5. The expression of CCR5 was increased in *myeKlf2^-/-^* mice compared to WT on both the days tested (n=5, p<0.05 for days 3 and 5 post CTX). **B.** Same strategy was followed and sorted by flow cytometry using antibody against CCR2 but there was no difference in expression between WT and *myeKlf2^-/-^* mice (p = insignificant). **C.** Representative flow cytometry analysis for gastrocnemius muscles that were obtained from WT and *myeKlf2^-/-^* mice at days 3 and 5 post CTX injury and stained and sorted using antibody against MerTK. The expression of CCR5 was increased in *myeKlf2^-/-^* mice compared to WT on both the days tested (n=5, p<0.01 for days 3 and 5 post CTX).

The number of cells expressing CCR5 is significantly greater in regenerating muscles of *myeKlf2^-/-^* compared to WT on day 3 post injury (73% in *myeKlf2^-/-^* vs. 61.2% in WT samples; n = 5; p< 0.05; Fig. 4A). This cell population remains elevated on day 5 in *myeKlf2^-/-^* samples (45.3% in *myeKlf2^-/-^* vs. 26.8% in WT, n = 5; p<0.05), but declines sharply after day 3 in WT.

KLF2 deficiency does not change the quantity of intramuscular CCR2-expressing cells on days 3 or 5 post injury (Fig. 4B). CCR2 is a receptor for the ligand CCL2, which is highly upregulated at early times in CTX-treated *myeKlf2^-/-^* muscle samples (Fig 1). This finding suggests that KLF2 regulation of cytokine receptor expression during muscle regeneration is receptor^-^specific.

We also compared expression of the C-mer receptor tyrosine kinase, MerTK, in regenerating WT and *myeKlf2^-/-^* muscle samples (Fig. 4C). MerTK defines a monocyte/macrophage phenotype that plays a role in resolving inflammation in other immune conditions [24, 25]. MerTK expression is elevated at day 3 in *myeKlf2^-/-^* compared to WT (17.7% vs. 10.9%, n = 5, p<0.01) and increases further by day 5 (24.5% in *myeKlf2*^-/-^ vs 6.6% in WT. Fig. 4C, n = 5, p<0.01).

### Markers of satellite cell activation and myogenesis appear earlier and are enhanced in regenerating muscles of myeKlf2^-/-^ mice

In addition to resolving inflammation, MerTK also helps initiate myogenesis. To examine whether the earlier resolution of inflammation in *myeKlf2*^-/-^ accelerates the progression of myogenesis, we measured expression of key markers of satellite cell activation, proliferation, and differentiation (Fig. 5). All satellite cells express Pax7, whereas only activated satellite cells express the transcription factors MyoD and Mrf6, which promote myogenesis. MyoD is one of the earliest markers of myogenic commitment. Myogenic regulatory factors (MRFs) are master regulators of skeletal myogenesis, among which Myf5 and Myf6 are the earliest expressed MRFs [26]. The appearance of these myogenic markers denotes the presence of activated satellite cells and newly forming fibers.

**Figure 5.**
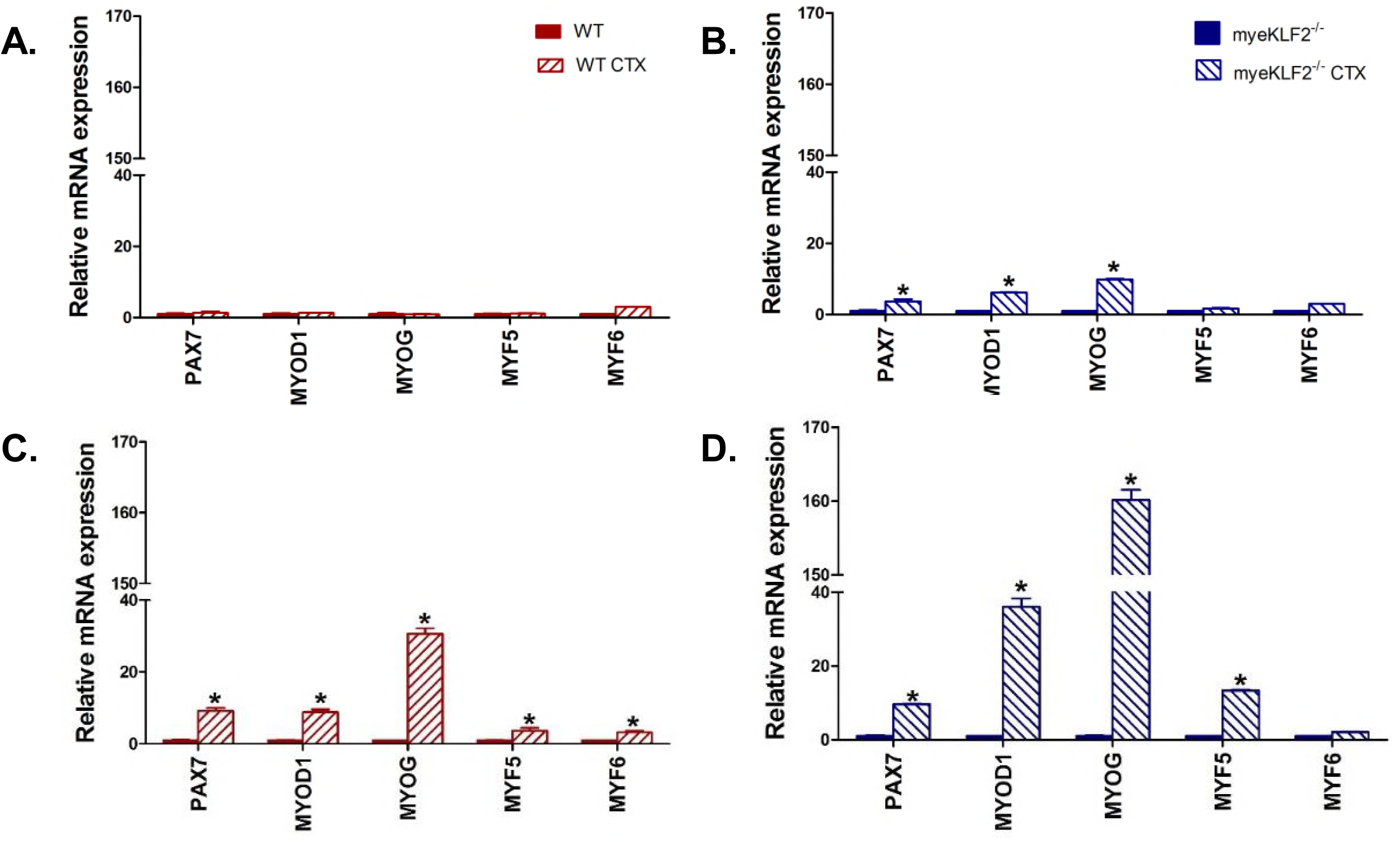
Markers of satellite cell activation and myogenesis appear earlier and are enhanced in regenerating muscles of *myeKlf2^-/-^* mice. **A & B.** RT-qPCR analysis for myogenesis markers in gastrocnemius muscles of WT (red) and *myeKlf2^-/-^* (blue) mice were taken at day 3 following placebo (solid fill) or CTX (striped) injury. **C & D.** RT-qPCR analysis for myogenesis markers in gastrocnemius muscles of WT (red) and *myeKlf2^-/-^* (blue) mice were taken at day 5 following placebo or CTX injury. ^*^ indicates significant difference at p < 0.05; n = 4.

Regenerating muscles of *myeKlf2^-/-^* mice express these markers earlier and at higher levels. Pax7, MyoD, and MyoG are already expressed at day 3 (Fig. 5B), are elevated 3- to 5-fold at day 5 (Fig. 5D), and remain elevated through day 8 (data not shown). Expression of Myf5 is increased by day 5 in *myeKlf2^-/-^* mice.

### Regeneration is advanced and completed normally in injured muscles of myeKlf2^-/-^ mice

To determine the consequences of these differences on the progression of muscle regeneration, we compared the histology of regenerating muscles of WT and *myeKlf2*^-/-^ mice (Fig. 6A). Uninjured muscles of both genotypes show phenotypically normal histology. By days 5 and 8 post injury, injured muscles of WT mice show extensive necrosis with abundant monocytes, evident on H & E stained samples as regions having mononuclear cells with blue stained nuclei. These histological indices coincide with the appearance and decline of the proinflammatory mediators and macrophages shown in Figs. 1 — 5. By day 14, small, newly forming myofibers with central nuclei appear. At day 21, some fibers show a differentiated adult phenotype with larger cross-sectional areas and peripheral nuclei.

**Figure 6.**
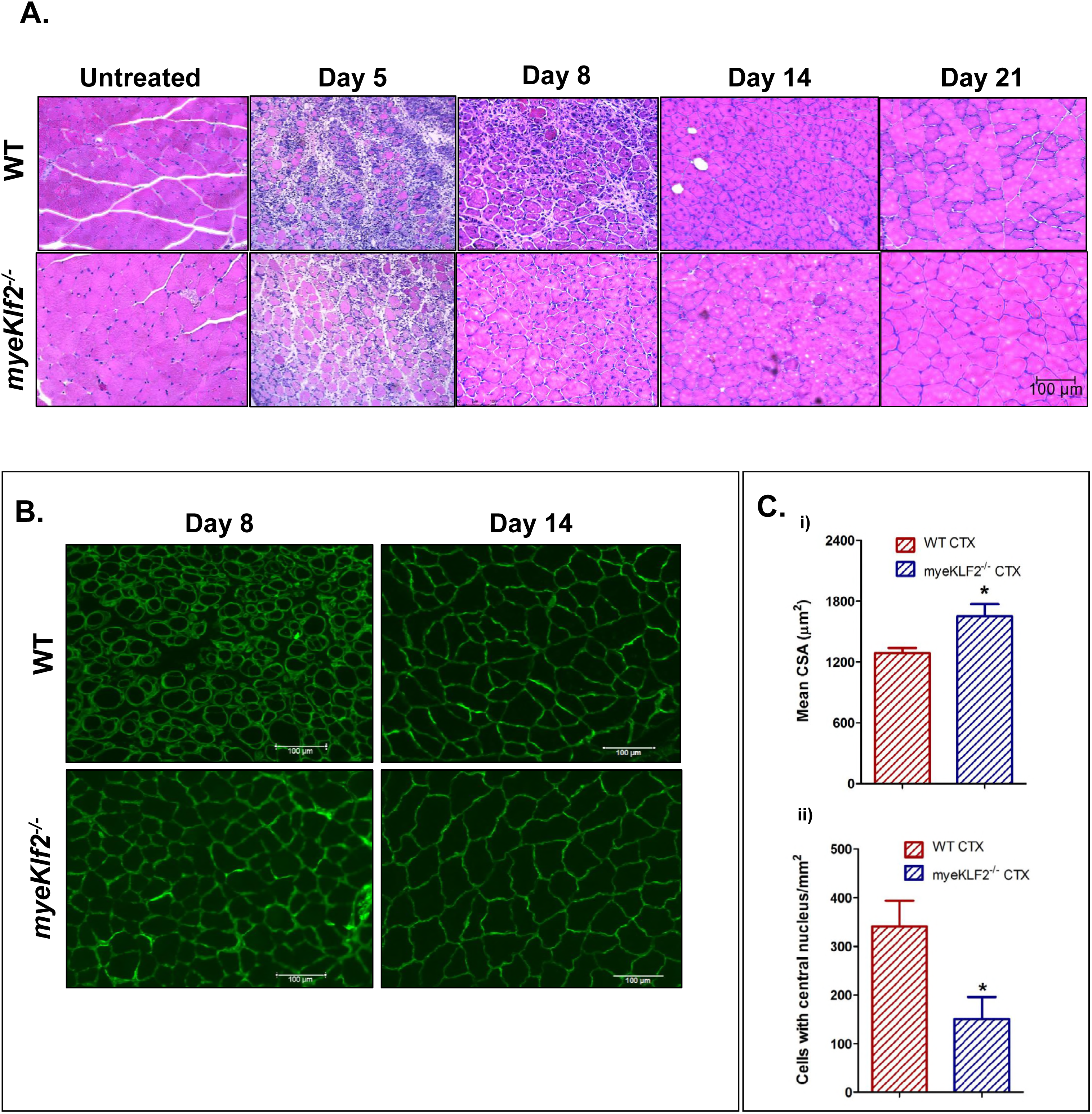
Regeneration is accelerated and completed normally in injured muscles of *myeKlf2^-/-^* mice. **A.** Representative H&E stained cross sections of gastrocnemius muscles from WT and *myeKlf2^-/-^* mice. Sections were taken from uninjured and CTX-injured muscles at days 5, 8, 14 and 21 post treatment. Scale bar 100 μm. **B.** Laminin labeled sections at days 8 and 14 post CTX treatment. Laminin was used to visualize fiber perimeters for determining cross-sectional areas. **C.** Mean cross-sectional areas and number of central nuclei in fibers of WT and *myeKlf2^-/-^* mice on day 14 after CTX injury. ^*^ indicates statistically significant difference at p<0.05 and n = 5 for each genotype.

Injured muscles of *myeKlf2*^-/-^ mice show an accelerated pattern of repair (Fig. 6B). Smaller necrotic regions with fewer monocytes are evident at day 5, consistent with an earlier resolution of inflammation. Newly forming fibers with small diameters and central nuclei are already present at day 8 and indicate that these muscles are transitioning to the myogenic phase of regeneration. By day 14, monocytes and necrotic areas have largely disappeared. Newly forming fibers have progressed to larger diameter, multi-nucleated adult phenotypes in which nuclei have moved to the fiber periphery (blue stained on H & E) with larger diameters. At day 21, muscles of *myeKlf2*^-/-^ mice show a regenerated adult architecture that is nearly indistinguishable from uninjured muscle.

To further quantify the progress of regeneration in WT and *myeKlf2*^-/-^ mice, we compared two indices of fiber maturation, mean cross-sectional areas and mean number of fibers with central nuclei at day 14 (Fig. 6B, 6C). Compared to WT, regenerating fibers of *myeKlf2^-/-^* show more fibers with larger cross-sectional areas and fewer central nuclei.

## Discussion

Acute skeletal muscle injury induces a robust immune response that is essential for the proper completion of repair [6]. This response follows a specific program of coordinated interactions between the immune system and the injured muscle.

Myeloid lineage cells play a central role in this process. Little is known about the mechanisms that determine myeloid lineage phenotypes and functions during skeletal muscle repair. This study examined the role of the transcription factor KLF2, a key determinant of myeloid cell activation state and phenotypes, in the immune response to skeletal muscle injury.

### Summary of Findings

Compared to WT, regenerating skeletal muscles of *myeKlf2*^-/-^ mice express 15- to 30-fold greater levels of inflammatory mediators known to play important roles in repairing injured muscle (*Cox2*, *IL-6*, *Ccl2*, *Ccl4*, *Ccl5*, and *Tnfα*). Their appearance coincides with the presence in the injured muscle of greater numbers of inflammatory Ly6C^+^ monocytes/macrophages that are phenotypically Ly6C^+^, CD11b^+^, F4/80^+^. The enhanced proinflammatory phase resolves earlier and progresses to an enhanced anti-inflammatory, regenerative phase. This transition is marked by a decline in Ly6C^+^ macrophages and their replacement by anti-inflammatory Ly6C^-^ macrophages, which are phenotypically Ly6C^-^, CD11b^+^, F4/80^+^. This transition occurs in both genotypes, but occurs earlier and more efficiently in regenerating muscles of *myeKlf2^-/-^* mice. Expression of CCR5, an inflammatory C-C motif receptor for (*Ccl4*) MIP1β and *Ccl5* (RANTES), is significantly elevated and persists past the initial inflammatory period, when it normally declines in WT. A unique MerTK^+^ cell population, not previously reported in regenerating skeletal muscles, increases in *myeKlf2^-/-^* mice during this transition. During the predominantly myogenic phase of repair, muscles of *myeKlf2^-/-^* show elevated levels of factors (MyoD1, MyoG, and Myf5) that promote the activation, growth, and differentiation of satellite cells into adult fiber types. In all phases of repair, the expected histological changes coincide with the appearance of pro- and anti-inflammatory mediators and cell populations. *myeKlf2^-/-^* muscles show larger necrotic regions with greater numbers of intramuscular monocytes in days 3 – 5 post injury, form new fibers earlier, and complete regeneration earlier with adult type fibers (day 14 vs. day 21). Collectively, these findings demonstrate that loss of myeloid KLF2 significantly enhances the initial proinflammatory immune response and, notably, that the enhanced inflammatory phase progresses to an earlier resolution of inflammation and an earlier completion of regeneration.

### Functions of myeloid KLF2 in the immune response to skeletal muscle injury

These findings indicate that the zinc-finger transcription factor, KLF2, either directly or indirectly, negatively regulates genes that determine myeloid cell phenotypes during skeletal muscle repair.

Injured skeletal muscles recruit mainly Ly6C^+^ monocytes from the circulation [5]. Ly6C^+^ monocytes are non-dividing, but mature within the injured muscle into macrophages capable of proliferation and differentiation [5]. If KLF2 negatively regulates genes which determine the Ly6C^+^ phenotype, its loss may bias their distribution and/or numbers in the circulation, making more available for recruitment. In addition, KLF2 may regulate genes that drive the *in situ* maturation of infiltrating Ly6C^+^ monocytes into macrophages. The increased abundance of Ly6C^+^, CD11b^+^, F4/80^+^ macrophages in injured muscles of *myeKlf2^-/-^* mice suggests that KLF2 regulates this transition, and that this phenotype plays an important role in the initial, inflammation phase of repair. In rheumatoid arthritis, reduction of KLF2 in hemizygous mice significantly increases the population of inflammatory Ly6C^+^, CD11b^+^, F4/80^+^ monocytes in sites of inflammation (10).

Myeloid-derived monocytes and macrophages produce the chemokines, cytokines and growth factors that are abundant in injured skeletal muscles and execute the various repair processes [27]. The significantly elevated expression of *Cox2*, *Ccl2*, *Ccl4*, *Ccl5*, *IL-6* and *Tnfα* in regenerating muscles of mye*Klf2*^-/-^ mice suggests that KLF2, either directly or indirectly, regulates their expression. Previous studies of injured muscle have shown that the cytokines TNFα and COX2 originate from both inflammatory macrophages and injured muscle fibers, and they participate in both inflammatory and regenerative functions [21, 28, 29]. A role of KLF2 in regulating their expression is consistent with previous reports from our laboratory and others that KLF2 functions as a central, negative regulator of these genes in other inflammatory conditions. KLF2 directly regulates *Ccl2*/MCP-1, *Tnfα*, and *Cox2* expression in various activated monocytes *in vitro* and *in vivo* [17]; and KLF2 deficiency in myeloid cells increases the expression of *Ccl2*/MCP-1, *Ccl5*, *IL-6* and *Cox2* in inflammatory, M1-type peritoneal macrophages [18].

IL-6 is secreted by inflammatory monocytes/macrophages and plays a central role in the inflammation phase of repair. The majority of the initial IL-6 pool is produced by monocytes recruited from the circulation [20]. IL-6 is also produced by activated satellite cells and this pool of IL-6 provides a key myokine for the formation and differentiation of new fibers [30]. Our finding that KLF2 negatively regulates IL-6 expression early in regenerating muscles is novel and may explain, in part, the ability of an enhanced inflammatory phase to positively drive the regenerative phase of repair. IL-6 produced by proinflammatory macrophages, together with other chemokines and factors, may promote activation of satellite cells and further raise IL-6 levels in the satellite cell niche, as proposed by Munoz-Canoves *et al*. [30].

The expression pattern of cytokine receptors on monocytes and macrophages determines their response to chemokines. The markedly enhanced expression of CCR5 and its persistent expression in *myeKlf2^-/-^* muscles suggests that KLF2 reversibly determines CCR5 expression in myeloid-derived intramuscular macrophages during muscle regeneration. Myeloid cells in *myeKlf2^-/-^* mice cannot elevate KLF2 and, consequently, cannot downregulate CCR5 expression, as occurs in T cells, monocytes, and myeloid-derived macrophages in other immune conditions [16, 17]. The enhanced and sustained expression of CCR5 may contribute to the larger population of Ly6C^+^ monocytes and Ly6C^+^ macrophages in injured muscle of *myeKlf2^-/-^* mice. In other immune cells and conditions, CCR5 receptors promote the migration of inflammatory monocytes to the site of injury and their differentiation into mature macrophages. For example, only activated T cells, after downregulating KLF2, express CCR5, and CCR5 is required for their proper migration [16]. CCR5 expression increases during differentiation of human monocytes and monocyte-derived macrophages [31, 32]. In atherosclerosis, the expression of CCR5 and CX3CR1 is required for Ly6C^+^ monocytes to accumulate in plaques [33].

The switch of macrophage phenotypes from Ly6C^+^ to Ly6C^-^ populations is required for regeneration to proceed [22, 34]. It is notable that the disappearance of Ly6C^+^ macrophages and their replacement with Ly6C^-^ macrophages persists in regenerating muscles of *myeKlf2*^-/-^ mice, indicating that myeloid KLF2 is not directly required for this transition. The transition may be driven by local signals related to phagocytosis, as suggested by Arnold et al [5] and as shown in other cell types [35-38].

The unique MerTK^+^ population of macrophages that increases during this transition in *myeKlf2^-/-^* muscles may contribute to resolving inflammation. MerTK is a member of the Tyro3/Axl/Mer (TAM) receptor tyrosine kinase family, whose functions in macrophages include phagocytosis of necrotic cells [39] and the subsequent dampening of inflammatory responses [40, 41]. After acute liver injury, an expanding population of monocyte-derived MerTK^+^ macrophages clears damaged cellular material and also secretes anti-inflammatory mediators that help resolve inflammation [24]. In wound healing, the population of Ly6C^-^, MerTK^+^, F4/80^+^ macrophages increases as inflammation resolves and regeneration proceeds [25]. In cardiac muscle damaged by ischemia, MerTK promotes the resolution of inflammation and its loss compromises repair [42, 43]. Our findings identify MerTK as a KLF2*-*regulated receptor that influences the transition from inflammation to regeneration during skeletal muscle repair.

### Key differences between myeloid KLF2 functions in skeletal muscle repair after acute injury, and its role in chronic inflammatory conditions

A more robust inflammatory phase of muscle regeneration in *myeKlf2*^-/-^ mice is consistent with known functions of myeloid KLF2 as a negative regulator of inflammatory genes and inflammatory immune cell profiles in other conditions [14-16]. Overexpression of KLF2 in leucocytes induces quiescence [44] and represses inflammatory genes [15]. In a mouse model of atherosclerosis, peritoneal macrophages of *myeKlf2*^-/-^ mice show a chronic inflammatory profile, produce elevated levels of cytokines, and show enhanced adhesion and infiltration into atherosclerotic lesions [18, 19].

On the other hand, our finding that the inflammation in injured muscles resolves in the complete absence of KLF2 was unexpected. In atherosclerosis and other chronic inflammatory conditions, a reduction in KLF2 levels in macrophages promotes inflammatory gene expression, and a subsequent rise in KLF2 terminates their expression. When KLF2 levels cannot rise, the inflammation is enhanced and persists, and tissue damage worsens [19]. A different myeloid cell program must operate during skeletal muscle repair. When inflammatory gene expression is enhanced by deleting KLF2, inflammatory Ly6C^+^ macrophages transition normally to an anti-inflammatory phenotype without the requirement for KLF2.

### Therapeutic implications

Skeletal muscle injuries related to aging, disease and sports are the most common musculo-skeletal conditions reported worldwide. Severely injured skeletal muscle has limited ability to repair and regenerate. This study suggests a novel strategy for improving outcomes after muscle injury. Conventional treatments for muscle injuries seek to limit inflammation. Our findings suggest that *enhancing a specific inflammatory program* by manipulating KLF2 in myeloid lineage cells can accelerate regeneration. Myeloid KLF2 is an attractive therapeutic target for several reasons. Targeting a transcription factor, rather than a single chemokine or receptor, provides an opportunity to controls a cluster of interacting genes that determine macrophage phenotypes and functions. Importantly, myeloid KLF2 is accessible from the circulation and controls an early step of repair that positively drives the overall progress of regeneration.

### Conclusions

The zinc-finger transcription factor, KLF2, plays a central role in muscle-immune cell interactions during skeletal muscle regeneration after injury. Loss of KLF2 in myeloid cells enhances the initial inflammation phase of repair and positively drives regeneration to a normal and earlier completion. This novel finding underscores the specificity of the immune response to different conditions, and identifies KLF2 as a central regulator of myeloid cell phenotypes and functions during skeletal muscle regeneration.

## Material and Methods

### Animals

*Klf2^fl/fl^* mice were generated and backcrossed to a C57BL/6 background as described [19]. *Klf2^fl/fl^* mice were crossed with transgenic mice expressing *Cre* recombinase under the control of the LysM promoter (LysM-*cre*, Jackson Laboratories; [45] to produce offspring with a targeted deletion of KLF2 in all myeloid-derived cells. *Klf2^fl/fl^* mice without the *Cre* recombinase transgene were used as controls. All mice were housed in pathogen-free conditions at the University of Cincinnati. All procedures and animal care were performed in accordance with the Guide for the Care and Use of Laboratory Animals (National Research Council of the National Academies, USA) and were approved by the Institutional Animal Care and Use Committee of the University of Cincinnati.

### Muscle Injury

Acute injury was produced by injecting Cardiotoxin (CTX; EMD Millipore, 100 μl of 10 mM stock dissolved in PBS) into the gastrocnemius muscle. CTX injury is an established model of skeletal muscle regeneration. It induces a reproducible sequence of changes that reflect the physiological repair process after traumatic injury [46]. The gastrocnemius muscle was surgically removed for analysis at time points up to 21 days post injury and the animals were sacrificed.

### RNA isolation and real^-^time quantitative PCR (RT-qPCR)

Total RNA was isolated from skeletal muscle and macrophages using mirVana miRNA isolation kit (Invitrogen). cDNA was synthesized using the Superscript IV reverse-transcription system (Invitrogen, # 18091050) and genomic DNA was eliminated using EZDNASE (Invitrogen, # 11766051). mRNA expression was quantified using gene-specific Taqman primers and probes (Applied Biosystems) following the manufacturer’s protocol, and normalized to 18sRNA levels. RT-qPCR was performed on an ABI7300 real^-^time PCR system (Applied Biosystems) using Taqman universal master mix (Applied Biosystems, #4440044). Relative gene expression was calculated using the ΔΔ*C_t_* method following the manufacturer’s instructions. All reactions were carried out in triplicate using RNA isolated from muscles of 4 - 8 mice at each time point.

### Histological analysis

Gastrocnemius muscles were removed, snap frozen in nitrogen-chilled isopentane, embedded in O.C.T. (Fisher Scientific) and kept at -80°C until use. Cryosections of 10μm thickness were prepared for haematoxylin/eosin (H&E) staining. Sections from the entire injured area were cut and analyzed quantitatively for the percentage of necrotic and regenerating myofibers, fiber cross-sectional areas, and the number of fibers with central nuclei. Sections obtained from at least three mice per time point were analyzed.

### Cell isolation and flow cytometry

Flow cytometry was performed using a suspension of single cells obtained from the muscle. Briefly, for cell isolation from tissue, injured muscles were placed in warmed DMEM (Life Technologies). The muscles were chopped into small pieces and enzymatic disaggregation was performed using a skeletal muscle dissociation kit (Miltenyi Biotec, #130-098-305). The dissociated sample was filtered through a 70 μm cell strainer (Fisher Brand). Red blood cells (RBCs) were lysed with RBC Lysing buffer (eBioscience, #00-4300-54) at room temperature for 10 min. After lysis, cells were centrifuged and the pellet was re^-^suspended in flow cytometry buffer (eBioscience, # 00-4222-26). Following the lysis of RBC cells the samples were blocked with CD16/32 for 10 min on ice. For flow cytometry analysis of monocytes and macrophages, the cells were stained with Ly6C-APC (eBioscience), CD11b-FITC (Biolegend) and Super Bright 436 (eBioscience), F4/80-PECy7 (Biolegend), MerTK-PerCP-eflour 710 (eBioscience), CCR5-APC Vio770 (Miltenyi Biotech), CCR2-PE (Biolegend). Sytox blue (Invitrogen) was used to exclude dead cells and debris, and unstained controls were used to set gates. Samples of 30,000 cells per run were analyzed in the flow cytometer (LSR Fortessa, BD Bioscience) and the resulting data was analyzed using FlowJo software (FlowJo, LLC). Only samples that passed the appropriate filters and quality checks were analyzed. Cell counts were expressed as the percentage of positive cells per sample. Cell suspensions were obtained from 4 — 8 mice for each time point studied.

### Statistical analysis

Statistical analysis was performed with GraphPad Instat program and values were expressed as mean ± SD. Multiple comparisons were tested by ANOVA, with Student Newman-Keuls post hoc analysis. A difference of *p* ≤ 0.05 was considered statistically significant.

## Acknowledgments

We thank Drs. Christina Wei and Lubov Timchenko, Cincinnati Children’s Hospital Medical Center (CCHMC) for assistance in developing the cardiotoxin mediated skeletal muscle injury model. We thank Dr. David Hui, University of Cincinnati for his critical comments on the manuscript. We thank the Flow Cytometry Core of CCHMC for the use of instrumentation and analysis software. Research reported in this publication was supported by the National Institute of Arthritis and Musculoskeletal and Skin Diseases of the National Institutes of Health under Award number RO1-AR063710 and by the National Institute of Heart Lung and Blood under award numbers R01HL130356 and R01HL105826. The content is solely the responsibility of the authors and does not necessarily represent the official views of the National Institutes of Health.

## Authors Contributions

PM, JBL, JH designed research

PM, TS, TLR performed research

PM, TS, TLR, SS, JBL, JH analyzed data and read manuscript

JBL contributed new reagents/analytic tools;

PM, JBL, JH wrote the paper

## Conflict of interest

The authors declare no conflict of interest.

**Abbreviations**
KLF2: Krüppel^-^like Factor 2
WT: wild type
*myeKlf2*^-/-^: mice with targeted knockout of KLF2 in all myeloid-derived cells
CTX: cardiotoxin

